# Common attentional mechanisms, but distinct computational dynamics, across Statistical and Reward Learning

**DOI:** 10.64898/2026.11.04.764241

**Authors:** Carola Dolci, Tom Verguts, Willem Jacob Louter, Elisa Santandrea, C. Nico Boehler

## Abstract

Past experience guides attention through statistical learning (SL) and reward learning (RL). Yet, whether they do so through similar neural and computational mechanisms remains unclear. In a first EEG visual-search experiment, we tested whether RL establishes latent spatial representations, previously demonstrated for SL. Afterwards, we directly compared RL and SL in a double-session experiment. Using multivariate decoding, Experiment 1 showed that high-reward locations could be decoded before target onset following a task-irrelevant, spatially-unspecific visual perturbation. Experiment 2 revealed that SL and RL established comparable neural representations, and similarly facilitated target selection and evidence accumulation (drift-rate). However, combining Rescorla-Wagner and Drift Diffusion Model revealed slower learning-rate for RL than SL. Pupillometry further tracked spatially selective expectation violations during SL, but reflected increased arousal during RL. Together, our results reveal how the brain routes distinct computational algorithms (probability estimation versus value caching) into a unified attentional priority map.

## Introduction

Adaptive behaviour depends on learning which information is likely to be behaviourally significant, using that knowledge to guide attention, thereby prioritising relevant information among competing sensory inputs^1,2^. Alongside goal-directed^3,4^ and stimulus-driven mechanisms^5,6^, recent work has established that attentional orienting is additionally guided by learning from past experience^7^. Yet, despite growing interest in experience-dependent attentional control, it remains unclear how different forms of experience establish attentional priorities, and whether they do so through similar or distinct processes.

Experience can guide attention through at least two distinct forms of learning. Statistical learning (SL) extracts frequency regularities in the environment, biasing attention towards locations where task-relevant information is more likely to occur^8,9^. Reward learning (RL), by contrast, assigns priority according to previous outcomes. Stimuli or locations associated with greater reward receive enhanced attentional priority, which can persist even when reward is no longer available or relevant for the current task^10,11^.

Despite their different core objectives (target prediction in the sense of probability estimation vs. reward caching in the sense of reinforcement history), in the field of attention, SL and RL have often been grouped under the broader category of experience-dependent mechanisms. Both emerge gradually through experience, influence selection relatively automatically, and operate with limited (or no) conscious awareness^12,13^. However, at the same time, theories of learning suggest fundamental differences between SL and RL. At the computational level, their distinction parallels the contrast between model-based and model-free computations, or between generative versus discriminative models^14^. Model-based/SL aims to learn the structure of the world. Based on accumulated regularities and without the need of explicit feedback, it creates internal representations of environmental contingencies to generate predictions of future events^15,16^. Model-free/RL, in contrast, aims to optimize behavioural outcomes. It arises from incremental value updating through trial-and-error experience^17,18^, where, based on feedback, it assigns different values to stimuli, actions or locations.

Importantly, evidence from computational neuroscience suggests that model-based and model-free learning rely on partially dissociable neural substrates, while their outputs can ultimately converge to guide behaviour^19^. This architecture is in line with theoretical accounts of attentional control in which multiple attentional control mechanisms, including SL and RL, converge within a common spatial priority map^20–22^. Consistent with this view, behavioural studies indicate that, within this map, SL and RL independently and additively shape attentional selection^23,24^. However, whether these behavioural similarities arise from common neural mechanisms remains largely unknown.

At the neural level most studies have focused on the process during target selection itself, showing that both forms of learning modulate multiple stages of target selection, from early attentional orienting to post-selection evaluation, as reflected in distinct event-related potential (ERP) components^25,26^. However, recent work has started to focus on how learning information is represented within the brain even in the absence of relevant sensory input, highlighting the importance of moving beyond studying stimulus-evoked activity. Specifically, perturbation-based electroencephalography (EEG) studies suggest that attentional priorities can be maintained in latent, activity-silent neural states, supported by transient changes in synaptic connectivity rather than by sustained neural firing, and revealed when the system is perturbed by a task-irrelevant sensory (“ping”) event^27,28^. Using such a visual ping, Duncan and colleagues (2023)^29^ showed that a high-frequency target location bias learned through SL could be decoded from the evoked EEG response before the search display appeared. This result suggested that statistical regularities can establish an anticipatory spatial representation that biases target selection in a way that can not be explained by active top-down expectations^30^. Whether RL establishes a comparable latent representation is unknown.

A complementary perspective comes from physiological measures. Pupillometry has long been linked to arousal and learning-related computations^31^, where studies revealed that reward expectancy is associated with enhanced pupil dilation, reflecting motivational engagement^32,33^. In contrast, the physiological signatures of SL remain less understood. Given that pupil responses are sensitive to violations of learned expectations^34^, they may also reflect violations of target predictions formed through SL. Comparing pupillary responses across SL and RL therefore provides a valuable opportunity to determine whether learning related to target probability and reward caching rely on common or distinct physiological mechanisms.

Here, across two experiments, we address these gaps by directly comparing SL and RL within a common experimental framework and tracking their convergence or divergence across multiple levels of analysis, from their neural representations and physiological responses to their computational dynamics. In Experiment 1 (Fig. 1a), we tested whether RL, like SL^29^, generates a spatial representation that can be maintained in a latent neural activity before target onset. Using a perturbation-based EEG approach, we tested whether information about a high-reward location could be reactivated by a task-irrelevant visual ping, while remaining undetectable during no-ping intervals.

**Figure 1.**
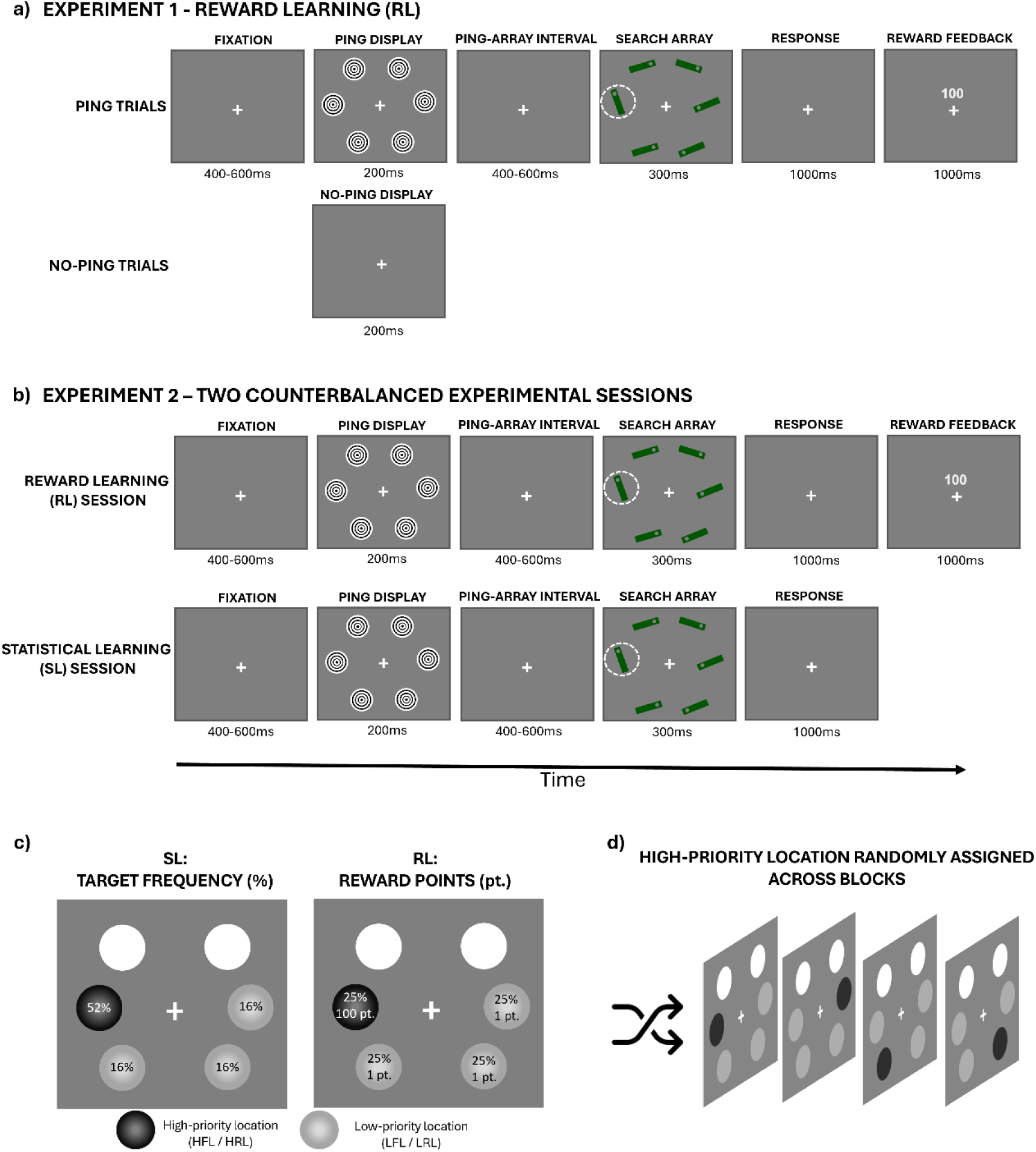
Experimental design. **a, b,** Example trial sequences for Experiment 1 and 2. In Experiment 1 (panel a), half of the trials contained the ping, whereas on the remaining no-ping trials, only the fixation cross was presented during the corresponding 200-ms interval. The target (circled for visualization only) was defined as the bar tilted ±25° from the vertical axis, whereas non-targets were tilted ±25° from the horizontal axis. Experiment 2 (panel b) consisted of two counterbalanced sessions, respectively focused on RL or SL. Here, each search display was always preceded by a task-irrelevant high-contrast ping stimulus. For RL (Experiment 1 and RL session in Experiment 2), correct responses were followed by reward feedback indicating either a high (100 points) or low (1 point) reward, according to the reward magnitude associated with the target location. When the response was incorrect the screen included only the fixation cross. **c,** Frequency/value distribution across locations during SL (left) and RL (right) session. In the SL session, targets appeared more often in one location (52%) and less often in the others (16%). In the RL session, targets appeared equally often in all the locations (25%), but one location yielded a high reward (100 points), contrasting with low reward in all other locations (1 point). **d,** Spatial distribution of high-priority (HFL/HRL) location across experimental blocks. The HFL/HRL was randomly rotated across the four possible target locations, such that each location served as the high-priority location three times across the 12 experimental blocks. Targets never appeared at the two upper display locations.

Having established this, Experiment 2 directly compared SL and RL within a common experimental framework (Fig. 1b). Using an identical task, participants completed two (counterbalanced) sessions in which attentional priorities were induced either by SL (high-vs. low-frequency target locations) or by RL (high vs. low reward locations). Combining multivariate EEG decoding, pupillometry, computational modelling, and ERP analyses, we provide a comprehensive characterization of SL and RL during the full cascade of target selection process, from anticipatory states until actual target selection and discrimination.

## Results

### Experiment 1: Reward learning facilitates target selection and establishes an anticipatory spatial representation at the latent neural network level

In order to confirm that we successfully induced a behavioural bias with the reward manipulation, accuracy and reaction times (RTs) were analysed using separate one-way repeated-measures ANOVAs with Reward Location (high vs. low) as a within-participant factor (Fig. 2a). As expected, participants were more accurate and responded faster when targets appeared at the high-compared with the low-reward locations [ACC: F(1,44) = 5.09, p = 0.02, η²_p_ = 0.10; RTs: F(1,44) = 16.21, p < 0.01, η²_p_ = 0.26].

**Figure 2.**
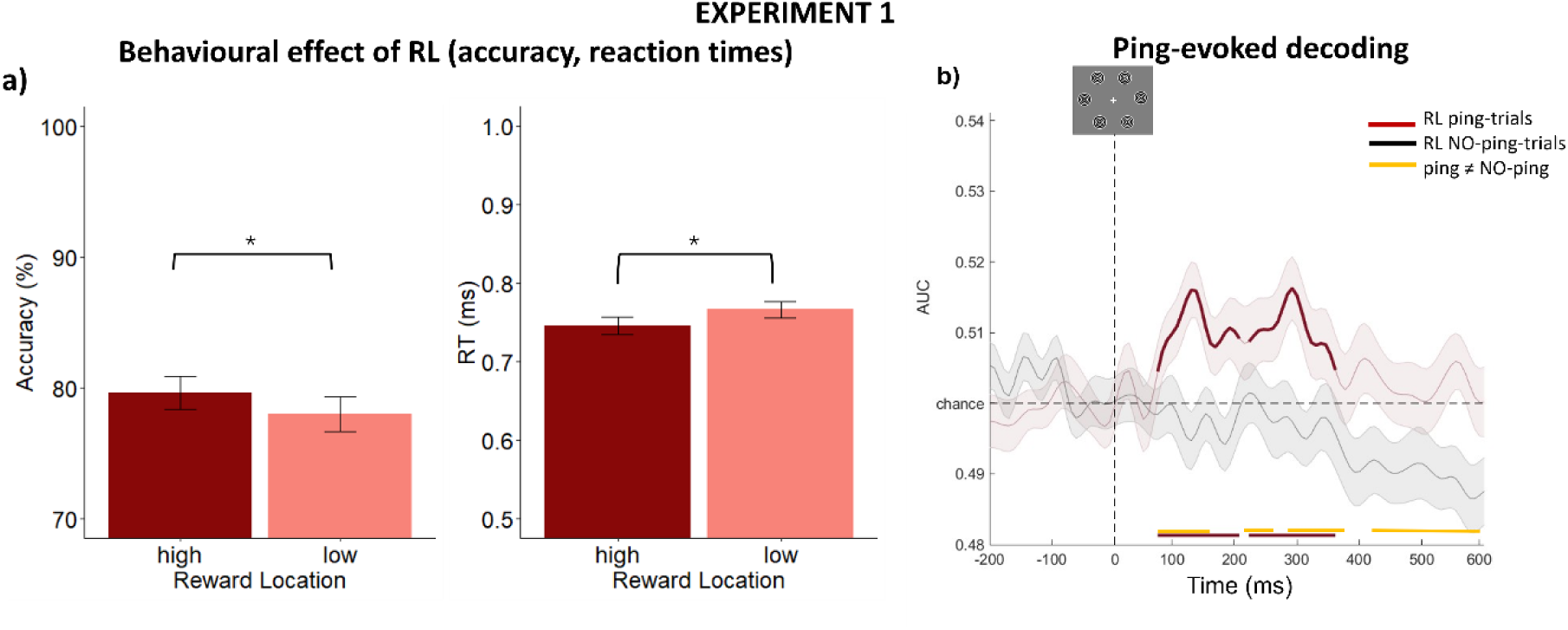
Behavioural effects and decoding results of RL. **a**, Mean accuracy (left) and reaction times (right) as a function of Reward Location (high vs. low). Error bars indicate s.e.m. across participants. Black asterisks indicate significant differences between reward conditions (p < 0.05). **b,** Time-resolved decoding accuracy for RL on ping (red) and no-ping trials (black). The thick line indicates above-chance-level decoding (0.5), and shaded areas indicate s.e.m. across participants. Horizontal bars above the x-axis indicate significant temporal clusters identified using cluster-based permutation tests (p < 0.05): within-condition clusters reflect decoding accuracy significantly above chance, whereas the yellow bar indicates a significant difference between ping and no-ping trials. Curves were smoothed for visualization using a 37-Hz low-pass filter.

We next asked whether high-reward location (HRL) information could be decoded from the EEG activity evoked by the task-irrelevant (and spatially non-selective) visual ping presented before the actual search array onset. Multivariate classifiers were applied separately to ping and no-ping trials during the 600-ms interval preceding the earliest possible search-array onset. Across experimental blocks, the HRL was rotated across the four possible target locations, which served as the four classification labels (Fig. 1d; see^29^ for similar approach). Critically, because the ping carried no information about the reward location, above-chance decoding from the ping-evoked activity would indicate that the perturbation revealed an existing reward-dependent spatial representation encoded in latent neural activity.

As shown in Figure 2b, results revealed significant (and cluster-corrected) decoding of the HRL already from 72 ms following ping onset (staying significant for roughly 300 ms). By contrast, HRL decoding did not significantly exceed chance during the corresponding interval on no-ping trials. Consistent with this difference, a direct two-sided cluster-based permutation test showed significantly greater decoding on ping than no-ping trials from 72 ms (remaining significant with some gaps until the end of the analysed time-window at 600 ms).

Together, these findings provide the first evidence that reward history not only facilitates target selection but also establishes neural trace that precedes the upcoming search display. This directly extends the anticipatory representation previously demonstrated for SL^29^, despite the fact that for RL, the expectation is about value conditional on a given location, not probability of target occurrence. This points to a common feature of experience-dependent attentional priority.

### Experiment 2 Behavioural analysis: Statistical and Reward learning similarly facilitate target selection

Demonstrating that both SL and RL generate spatial representations before search onset does not necessarily imply that they rely on the same underlying mechanisms or exhibit similar learning dynamics during all the stages of target selection. To address this question, Experiment 2 employed a multimodal approach to directly compare SL and RL across anticipatory, target-selection, and post-selection stages of attentional processing within a counterbalanced two-session, within-participant design using the same visual-search task as in Experiment 1 (but including ping trials only).

In terms of behaviour, accuracy and RTs were analysed using 2×2 rm-ANOVAs with Learning Paradigm (SL, RL) and Location (high, low) as within-participant factors (Fig. 3a). High and low locations corresponded to the high- and low-frequency target locations (HFL and LFL) in the SL session and to the high- and low-reward locations (HRL and LRL) in the RL session. Performance was faster and more accurate at high-than low-priority locations, as reflected by a significant main effect of Location [accuracy: F(1,29) = 31.02, p < 0.01, η^2^_p_ = 0.51; RTs: F(1,29) = 52.85, p < 0.01, η^2^_p_ = 0.64]. There was no main effect of Learning Paradigm [accuracy: F(1,29) = 0.20, p = 0.65; RTs: F(1,29) < 0.01, p = 0.99], indicating that, overall, performance did not differ between the two experimental sessions. However, a significant interaction emerged between Learning Paradigm and Location [accuracy: F(1,29) = 4.69, p = 0.03, η^2^_p_ = 0.13; RTs: F(1,29) = 11.55, p < 0.01, η^2^_p_ = 0.28], indicating a larger benefit for SL (accuracy: 3.61%; RTs: 31.5 ms) than RL (accuracy: 1.53%; RTs: 10.6 ms).

**Figure 3.**
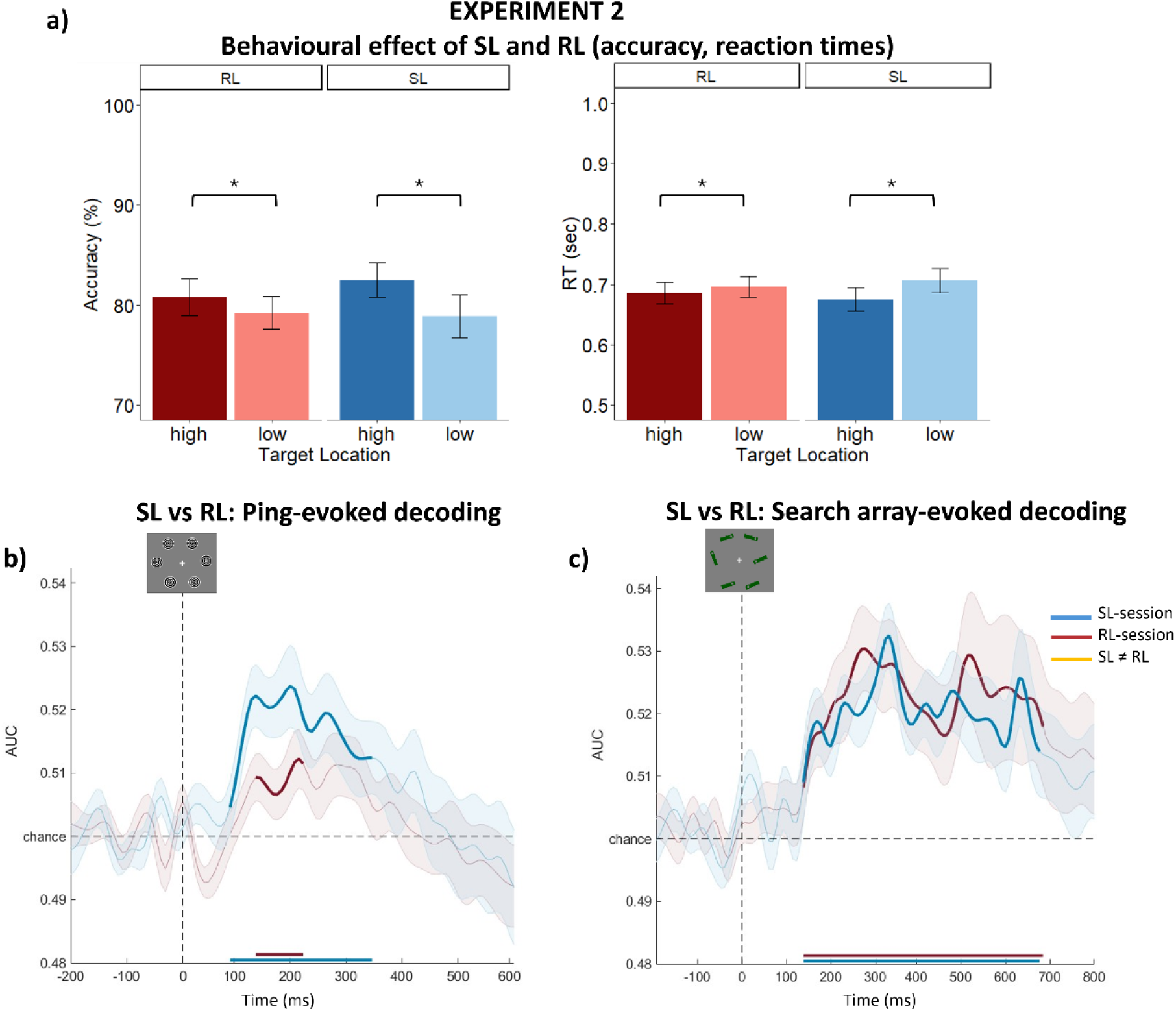
Behavioural data and decoding of SL and RL. **a**, Mean accuracy (left) and reaction times (right) as a function of learning paradigm (RL vs. SL) and Location (high vs. low). Error bars represent the standard error of the mean. Black asterisks indicate significant differences between conditions (p < 0.05). **b,c,** Decoding accuracy time-locked to the onset of the task-irrelevant ping display (b) and search array (c), shown separately for SL (blue) and RL (red). In all panels, chance-level decoding corresponds to 0.5. Horizontal bars above the x-axis indicate significant temporal clusters identified using cluster-based permutation tests (p < 0.05) against chance. In neither case did we observe any differences between SL and RL (as indicated by the absence of yellow lines marking significant temporal clusters). Shaded areas represent the s.e.m. across participants. Curves were smoothed for visualization using a 37-Hz low-pass filter.

This difference could partly reflect inter-trial priming, which is a potentially independent contributor to standard SL effects^9,29^. Because targets occurred more frequently at the HFL during SL, consecutive presentations at the same location were more frequent than during RL, where target location was equiprobable. We therefore repeated the analysis after excluding trials in which the target appeared at the same location as on the preceding trial (34% of trials in the SL session and 24.6% in the RL session). This analysis confirmed a significant main effect of Location, revealing better performance for the high (vs. low) location [accuracy: F(1,29) = 23.44, p < 0.01, η^2^_p_ = 0.44; RTs: F(1,29) = 18.47, p < 0.01, η^2^_p_ = 0.38] and the absence of a main effect of Learning Paradigm [accuracy: F(1,29) = 0.03, p = 0.85; RTs: F(1,29) = 0.04, p = 0.82], further supporting the comparability of the RL and SL sessions. Importantly, no significant interaction emerged between the two factors [accuracy: F(1,29) = 1.94, p = 0.17; RTs: F(1,29) = 0.87, p = 0.35], demonstrating that, after controlling for inter-trial priming, SL (accuracy: 2.59 %; RTs: 15.7 ms) and RL (accuracy: 1.40%; RTs: 10.5 ms) produced comparable effects.

### Experiment 2 Decoding: Statistical and Reward learning similarly shape latent anticipatory and active target representations

We next tested whether this behavioural similarity extended to their neural representation. Using the same decoding approach as for Experiment 1, we first compared anticipatory latent activity. Separate multivariate classifiers were applied to ping-evoked EEG activity during the SL and RL sessions to decode the HFL and HRLs, respectively.

Time-resolved cluster-based permutation tests against chance (0.5) revealed significant HFL decoding from 88 to 346 ms after ping onset, replicating previous evidence^29^. The HRL was similarly decoded from 135 to 221 ms, confirming the RL effect observed in Experiment 1 (Fig. 3b). Although SL decoding was qualitatively stronger and more sustained, a direct two-tailed cluster-based permutation test revealed no significant difference with RL.

Next, we tested whether this similarity between SL and RL persisted once the search array appeared and target selection began. Separate classifiers were thus applied to decode the HFL and HRL from search- array-evoked EEG activity. Importantly, to ensure that classification reflected the learned attentional priorities rather than the physical location or features of the current target, actual target locations and target features were balanced within each decoding class (see Methods). Thus, above-chance decoding indexed information associated with the learned HFL or HRL independently of where the target appeared on a given trial. Importantly, similarly to what has been done for behavioural analysis, these results remained qualitatively unchanged after controlling for inter-trial priming (see Supplementary Materials).

As shown in Figure 3c, time-resolved cluster-based permutation tests against chance revealed that neural activity related to the respective high-priority location during search could be decoded from 139 to 678 ms for SL, and from 139 to 686 ms for RL. A direct two-tailed cluster-based comparison again revealed no significant difference between SL and RL, confirming very similar dynamics.

Overall, EEG activity hence suggests a closely matched neural dynamic of SL and RL, from latent spatial representations before target onset to their expression during subsequent selection. This convergence extended to ERP activity, with only a minor divergence in the Sustained Posterior Contralateral Negativity (SPCN) component (see Supplementary Materials).

### Experiment 2 Pupillometry: Reward learning enhances overall pupil responses, whereas statistical learning tracks spatial expectations

Despite their similar neural dynamics, SL and RL may differ in the physiological states accompanying task execution and learned spatial priorities. We therefore examined pupil responses during target selection.

We first tested whether the two sessions differed in overall tonic pupil size independently of search- evoked responses. Absolute pupil diameter measured in the time-window from -200 to ping-display onset was analysed using a one-way rm-ANOVA with Learning Paradigm (RL, SL) as a factor. No significant difference was found [F(1,28) < 0.01, p = 0.97], indicating comparable baseline pupil size across the two experimental sessions.

We then analysed baseline-corrected task-evoked pupil responses between 800 and 1900 ms following search-array onset using a 2×2 rm-ANOVA with Learning Paradigm (SL, RL) and Location (high, low) as within-subject factors. Results showed a significant main effect of Learning Paradigm [F(1,28) = 30.86, p< 0.01, η^2^_p_ = 0.52] with larger pupil dilation in response to the search displays for RL (vs. SL), suggesting greater arousal during this session. Furthermore, we did not observe a main effect of Location [F(1,28) = 2.43, p= 0.13], but we did find a significant interaction between the two factors [F(1,28) = 13.70, p< 0.01, η^2^_p_ = 0.32]. Post-hoc t-tests (two-tailed) showed a significant difference between the high (vs. low) Location, but only during the SL session [t(28) = 3.93; p < 0.01, Cohen’s d = 0.13] and not in the RL session [t(28) = 1.53; p = 0.13] (Fig. 4a). Again, after controlling for inter-trial priming effect, the general pattern of results remained the same (see Supplementary Materials).

**Figure 4.**
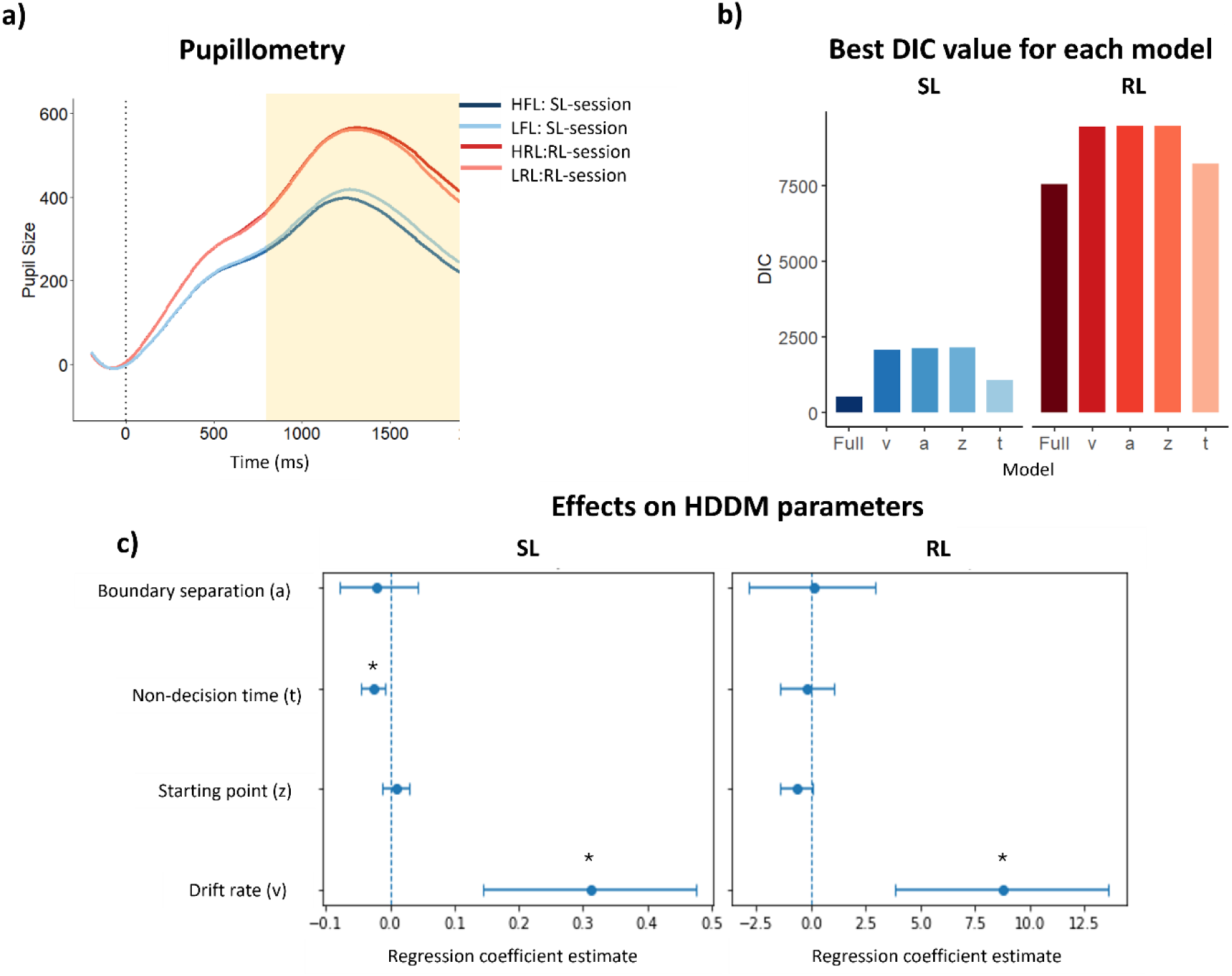
Pupil responses and computational modelling. **a**, Mean pupil diameter time-locked to search- array onset for high and low conditions during the SL (blue) and RL (red) sessions. The yellow shaded region indicates the analysis window (800–1900 ms) used for the statistical analysis of pupil responses. **b**, best DIC values for each of the five tested HDDMs across the different learning rates. The plot on the left represents the SL session and on the right the RL session. Note that the absolute DIC values should not be compared across SL and RL, as the Q-difference was generated from independently fitted models. **c**, Posterior estimates of the regression coefficients linking the trial-wise Q-difference of the four DDM parameters in the SL (left) and RL (right) sessions. Points indicate posterior means and horizontal bars show 95% credible intervals. Positive coefficients indicate that larger learning-related differences were associated with increases in the corresponding parameter, whereas negative coefficients indicate decreases. Credible intervals that do not include zero provide evidence for a reliable modulation (indicated with *). The vertical dashed line indicates a regression coefficient of zero.

Thus, pupil responses revealed a dissociation that was not apparent in the neural decoding results, suggesting that SL and RL triggered different internal physiological states. RL elicited a larger overall task- evoked response, consistent with greater motivational engagement or arousal (without this leading to a generally larger pupil size already at baseline), but showed no reliable modulation by reward location. SL, in contrast, produced a spatially selective pupil response, with greater dilation for targets appearing at low- than high-frequency locations, consistent with the established sensitivity of pupil dilation to surprise and violations of learned expectations.

### Experiment 2 Computational modelling: Statistical and reward learning converge on evidence accumulation but differ in their learning rate

We next turned to computational modelling to quantify how quickly SL and RL update attentional priorities across learning sessions and how these learned signals are translated into behavioural decisions. Specifically, we combined a Rescorla-Wagner (RW) model^35^ with a hierarchical drift diffusion model (DDM)^36^. The RW model was used to estimate a trial-wise learning signal reflecting the evolving value of each target location. Specifically, for each trial we computed a variable representing the difference between the estimated value of the current target location and the average value of the remaining locations, such that increasing values indicated a stronger learned preference for the high-frequency target location (in SL session) and high- reward (in RL session) location (henceforth, Q-difference). The resulting trial-wise Q-difference values were entered as regressors into separate HDDMs for the SL and RL sessions, allowing them to modulate different combinations of the diffusion-model parameters, namely drift rate (v); starting point (z); non-decision time (t); and boundary separation (a). Because it is unclear how quickly SL and RL were acquired, Q-difference was calculated using different learning rates, relying on subsequent model selection based on DIC values (see the Supplementary Material for an alternative updating algorithm that provided a poorer fit to the data).

Model comparison indicated that, in both sessions, the best-fitting model allowed the trial-wise Q- difference to modulate all four diffusion parameters simultaneously. Importantly, the optimal learning rate differed markedly between the two tasks. In the SL session, the best-fitting model was obtained with a relatively high learning rate (α = 0.40), whereas in the RL session the best fit was achieved with a much lower learning rate (α = 0.001), suggesting that spatial priorities were updated more rapidly during SL than during RL (Fig. 4b).

In the SL session (Fig. 4c), higher Q-difference values were associated with a significant increase in drift rate (v; posterior mean = 0.31, 95% credible interval (CrI) = [0.14, 0.47]), indicating faster accumulation of sensory evidence for targets appearing at the learned HFL. Higher Q-difference values were also associated with a small reduction in non-decision time (t; posterior mean = −0.03, 95% CrI = [−0.04, −0.01]), consistent with faster perceptual and motor processing (although this effect was no longer significant after controlling for inter-trial priming repetition, see Supplementary Materials). In contrast, there was no evidence that the Q-difference signal influenced the starting point (z; posterior mean = 0.01, 95% CrI = [−0.01, 0.03]) or boundary separation (a; posterior mean = −0.02, 95% CrI = [−0.07, 0.04]).

In the RL session, the Q-difference signal was also associated with a positive modulation of drift rate (v; posterior mean = 8.78, 95% CrI = [3.84, 13.60]), indicating enhanced evidence accumulation for targets at the HRL. By contrast, there was no evidence that the Q-difference signal influenced starting point (z; posterior mean = −0.66, 95% CrI = [-1.41, 0.01]), non-decision time (t; posterior mean = −0.17, 95% CrI = [−1.42, 1.06]) or boundary separation (a; posterior mean = 0.16, 95% CrI = [-2.82, 2.91]) (Fig. 4c).

## General Discussion

Attention is shaped by learning both what is likely and what is valuable^13,23^. Across two experiments, we show that SL and RL produced remarkably similar behavioural benefits, neural representations and convergent effects on evidence accumulation (drift rate) during decision making. At the same time, they clearly differed in their physiological responses and the rate of information acquisition, suggesting mechanistic convergence through partly dissociable process dynamics.

More specifically, a first major finding is that the maintenance of attentional priorities in latent neural states is not unique to SL. Previous work showed that a high-frequency target location acquired through SL can be recovered from the EEG response evoked by a task-irrelevant visual perturbation before search onset, despite not being detectable in ongoing activity^29^. Experiment 1 convincingly demonstrated that this principle extends to RL, despite the fact that the learned information is fundamentally different. The present results therefore suggest that also for RL, a latent network representation develops, possibly in the sense of changed synaptic weights along the visual hierarchy, which can only be made visible by active perturbation. Experiment 2 replicated this spatial representation of RL and directly compared it with SL. Both high-frequency and high-reward locations could be decoded before target onset, with no reliable difference in decoding strength between learning conditions.

The latent anticipatory representation observed during RL is particularly informative because, unlike target frequency, reward history provided no information about future events. In line with the predictive role of SL^14^, maintaining information about the high-frequency target location is directly beneficial for behaviour and it allows attentional resources to be, potentially, allocated in advance^37,38^. Such a predictive value concerning target locations was absent in our RL manipulation. In our task, reward acted exclusively as feedback contingent on target location, which was determined beyond participants’ control. Participants could neither predict nor influence where the next target would appear and prioritising the high-reward location could not increase the probability of finding the target and, consequently, the likelihood of getting a higher reward. Rather, participants could only cache reward values, which is relevant only in a conditional sense if the target was at that location. Nevertheless, information about the high-reward location was preserved in latent neural activity and reactivated even before search onset, suggesting the presence of a unique mechanism that can preserve learned information that serves for distinct functions, i.e., predictions about future events in SL and value caching in RL. Importantly, the modelling results indicated that the benefit was still exclusively related to an increase in drift rate, much like for SL, suggesting that in both cases, the actual process of evidence accumulation was improved.

At the active search stage, convergence between the neural representations of SL and RL became even more pronounced. Following search onset, high-frequency and high-reward locations could reliably be decoded over almost identical temporal intervals, again with no reliable difference between their time courses. ERP results related to attentional selection and target processing also largely converged (see Supplementary Materials), as did behaviour. Targets were selected more efficiently at both high-frequency and high-reward locations and, after controlling for immediate location repetitions, the magnitude of this spatial benefit was comparable between SL and RL. Computational modelling further formalized this convergence pattern. Once trial-by-trial learning (Q-difference) had been estimated, both SL and RL modulated the drift rate HDDM parameter, showing that stronger learned priority was associated with faster accumulation of task-relevant evidence towards a decision. Thus, despite originating from different sources of experience, both forms of learning ultimately produce comparable spatial priorities that translate similarly into behavioural decision making.

Importantly, the pupillometry findings suggest that SL and RL preserve distinct processing signatures despite the many elements of convergence. Pupil size, which we simultaneously recorded here, is known to be reflective of different relevant concepts, including surprise following violations of expectation^34^, arousal^39^, and cognitive effort or motivational engagement^32,33^. It therefore provides a valuable window into aspects of learning that may remain inaccessible to behavioural measures or neural decoding alone. During SL, task-related pupil dilation was larger when targets appeared at low- than high-frequency locations. This spatially selective response is consistent with evidence linking pupil dilation to surprise and prediction-error signalling^34,40,41^. Because each target occurrence provides information about the spatial structure of the environment, targets appearing at low-frequency locations violate the learned probability distribution and generate a greater mismatch between prediction and observed events. In contrast, RL produced a qualitatively different task-related pupil pattern. Pupil dilation was greater overall during RL than SL, consistent with enhanced arousal or motivational engagement when performance carries monetary consequences^31,42^. Critically, pupil responses did not reliably distinguish between high- and low-reward locations. Selectivity cannot be attributed to a failure to learn reward information. The high-reward location robustly facilitated behaviour and was represented neurally before and during the search task. Rather, the pupillary dissociation is consistent with RL relying on different processes than predictive SL. RL assigns and maintains values associated with individual locations without generating prediction about target appearance. Accordingly, unlike low-frequency location in SL, a target appearing at a low-reward location was not a violation of an expectation. Consistent with this account, SL, but not RL, showed a modest modulation of the non-decision time HDDM parameter, suggesting an additional influence on processes outside evidence accumulation^43^, although this effect did not remain robust after controlling for inter-trial repetitions (see Supplementary Materials).

Consistent with these different learning computations, i.e., prediction versus value caching, the dynamics through which spatial priorities were acquired also differed substantially. Trial-wise estimates were updated more rapidly during SL than RL. During SL, each target occurrence directly provided evidence about the spatial distribution of future targets and could therefore immediately update the current spatial prediction. During RL, by contrast, a target occurrence alone provided no information about spatial value. Value could only be established by linking the target location to the reward outcome that followed. Building a stable representation of spatial value therefore required integrating location–feedback associations across experiences. Consistent with this account, visual attention studies have shown that spatial frequency information is learned and exploited more efficiently than spatial reward magnitude information^44,45^. These findings thus suggest that there is no single temporal profile of experience-dependent mechanism. Rather, the rate at which attentional priorities emerge depends on the information provided.

Although our paradigm was not designed to distinguish model-based from model-free learning, this theoretical distinction provides an interesting parallel to the SL and RL processes investigated here. Model- based learning relies on an internal representation of environmental contingencies to prospectively evaluate future states and outcomes, conceptually resembling the predictive function of SL in our task. Model-free learning, in contrast, acquires values directly through trial-by-trial feedback, without requiring prospective knowledge of future states^17,18^, paralleling the value-based representation acquired through RL. Both models are simultaneously present in the brain and, although through partially dissociable neural substrates, they ultimately converge to guide behaviour^19^. Consistent with this possibility, studies manipulating SL and RL within a common attentional task suggest that the two sources of experience can make separable contributions to the same spatial priority map. Independent and additive influences of SL and RL have been reported behaviourally^24^, in oculomotor selection^23^, and in electrophysiological markers of attentional selection^46^. In our study we extended these investigations by identifying both points of convergence and divergence between SL and RL during the whole process of target selection. The resulting picture suggests that SL and RL ultimately give rise to remarkably similar attentional priority representations, which can be maintained in latent neural states and subsequently expressed during target selection, where they similarly facilitate performance and evidence accumulation. At the same time, differences in physiological responses and learning rate indicate that these representations may maintain their distinct functional roles.

More broadly, our findings establish a link between computational accounts of learning and theories of visual attention by showing how different forms of experience are transformed into attentional priorities. Despite encoding fundamentally different information, SL and RL converged onto a common experience- dependent mechanism, that preserved distinct functional signatures reflecting their different functional roles. SL contributes to attentional control by generating predictive representations of future events, whereas RL contributes by maintaining representations of learned value derived from past outcomes.

## Materials and Methods

The present study was conducted in accordance with the Declaration of Helsinki and approved by the ethical committee of the Faculty of Psychology and Educational Sciences of Ghent University (code 2025/11).

### Participants

Sample sizes were determined a priori based on previous studies using the visual-ping approach during SL^29^. Given that the effect of RL on ping-evoked spatial representations was unknown, and that previous work has suggested that spatial attentional biases might be weaker in RL than SL^47^, we recruited approximately twice the number of participants used in previous pinging studies, resulting in a final sample of 45 healthy volunteers in Experiment 1 (20 males; mean age = 23.8 years, SD = 3.55). In Experiment 2, where we included only ping trials, we recruited 30 participants (9 males; mean age = 24.1 years, SD = 2.90). All participants had normal or corrected-to-normal vision, were naïve to the purpose of the study, and had not previously participated in related experiments.

Participants received monetary compensation. In Experiment 1 and the RL session of Experiment 2, compensation consisted of a base payment of €30 plus a performance-dependent bonus of up to €8. For the SL session of Experiment 2, participants received €28. All participants provided written informed consent prior to participation.

### Apparatus and stimuli

Both experiments used the same visual-search paradigm. Participants were seated in a dimly lit, electromagnetically shielded room, 97 cm from a BenQ XL2420TE LCD monitor (1920 × 1080 pixels; 240- Hz refresh rate), using an adjustable chinrest. Pupil diameter was recorded using an EyeLink 1000 Plus desktop-mounted eye tracker positioned between the participant and the monitor (participant-to-eyetracker distance was 52 cm). The experiment was programmed and presented using PsychoPy (v1.84.2)^48^. Data quality and participants’ general task and central-fixation compliance were monitored remotely by the experimenter from an adjacent room.

The visual-search task was adapted from paradigms previously shown to elicit experience-dependent attentional biases^49,50^. Each search display consisted of six rectangular bars (2.0° × 0.5° of visual angle), arranged across an invisible circle 6° from a central white fixation cross (0.2° × 0.2°; RGB: 255, 255, 255; luminance: 190.2 cd/m²), which participants were asked to fixate continuously, and presented on a grey background (RGB: 66, 66, 66; luminance: 11 cd/m²). Equiluminant bars appeared in either green (RGB coordinates: 0, 76, 0; luminance: 11 cd/m2) or red (RGB values: 132, 0, 0; luminance: 11 cd/m2). Two bars were located in the upper visual field, two along the horizontal meridian, and two in the lower visual field. Each bar contained a small gap of the same grey colour as the background and positioned at either the top or bottom edge. The target was defined as a bar tilted ±25° from vertical, while distractors were tilted ±25° from horizontal. On each trial, participants were required to identify the position of the gap (upper or lower) within the target item. They responded by pressing the “Z” key with their left index finger if the gap was in the upper part, and the “M” key with their right index finger if it was in the lower part. Importantly, as in previous work^50^, the target could appear at four locations along the horizontal meridian or in the lower visual field, but never at the two upper locations, which contained filler items only. This restriction was introduced because the N2pc, one of the ERP components of interest, is typically attenuated for stimuli presented in the upper compared with the lower visual field^51,52^.

### General trial procedure

Unless otherwise specified below, trial structure was identical across experiments and learning conditions. Each trial began with a fixation display lasting 400–600 ms. On trials containing a perturbation stimulus, a task-irrelevant high-contrast “ping” was subsequently presented for 200 ms. The ping consisted of six concentric circular stimuli (2° diameter), each comprising an outer ring containing five smaller nested black- and-white circles, positioned at the same six locations as the subsequent search items. Following a further 400–600-ms interval, the search array was presented for 300 ms, followed by a fixation display for 1000 ms. Participants could respond from search-array onset until the end of this interval, yielding a total response window of 1300 ms. In RL blocks, a reward feedback was presented at the end of the trial: following correct responses, either “100” or “001” points (RGB: 255, 255, 255; luminance: 190.2 cd/m²) were displayed above the fixation point. No feedback was provided following incorrect responses or during SL session. Participants were instructed to maintain central fixation throughout the task (see Fig. 1).

### Experiment 1: Reward learning and perturbation manipulation

Experiment 1 comprised a single RL session of 1248 trials divided into 12 blocks of 104 trials. Target occurrence was spatially uniform: the target appeared equally often at each of the four possible target locations (25% each). Reward magnitude, however, varied systematically across locations. Within each block, one location was designated as the high-reward location (HRL), whereas the remaining three served as low-reward locations (LRLs). Correct responses to targets at the HRL yielded 100 points, whereas correct responses at LRLs yielded 1 point. At the end of the experiment, the accumulated points were converted into monetary rewards, with a maximum possible bonus of €8 which was added to the predetermined base payment (which participants were informed about before the experiment). Specifically, correct responses at the HRL (i.e. 100 points) yielded €0.02, whereas correct responses at LRLs (i.e. 001 point) yielded €0.002.

The HRL was systematically rotated across blocks such that each of the four possible target locations served as the HRL three times over the course of the experiment (see Fig. 1d). This was done in order to make the manipulations less obvious (which is particularly important to keep RL not explicit in Experiment 2) and to trigger a new learning process regularly. Additionally, this procedure is important in order to be able to collapse EEG data across hemispheres, thus minimizing noise due to inter-individual variability in brain asymmetries^53^.

Critically, Experiment 1 additionally manipulated the presence of the task-irrelevant ping. In each experimental block, half of the trials contained the ping display described above, whereas the remaining half were no-ping trials. The trial sequence and timing were identical across conditions; on no-ping trials, however, only the fixation cross was presented during the interval corresponding to the ping display. Ping and no-ping trials were randomly intermixed within each block.

### Experiment 2: Statistical versus Reward learning

Experiment 2 comprised two sessions, one manipulating SL and the other RL. Sessions were separated by 7 ± 1 days, and their order was counterbalanced across participants. Each session contained 1200 trials divided into 12 blocks of 100 trials. Participants completed a 16-trial practice phase before each experimental session.

The RL session followed the same reward-learning manipulation as Experiment 1. Unlike Experiment 1, however, every trial in Experiment 2 contained a ping stimulus before search-array onset. This design was motivated by the findings of Experiment 1, in which reliable anticipatory decoding of RL emerged only on ping trials, together with previous work showing stronger SL effect following ping- (vs. no- ping) evoked decoding^29^.

In the SL session, the spatial probability of target occurrence was manipulated. Within each block, one location was designated as the high-frequency target location (HFL), containing the target on 52% of trials. Each of the remaining three low-frequency target locations (LFLs) contained the target on 16% of trials (Fig. 1b). As for the HRL, the HFL was systematically rotated across blocks such that each of the four possible target locations served as the high-priority location three times during the experiment (Fig. 1d). As in the RL session, every trial contained the task-irrelevant ping before the search display, enabling the reactivation and decoding of latent spatial representations associated with the statistically high-priority locations.

### Electrophysiological recording and pre-processing

EEG data were recorded using a Biosemi ActiveTwo system using a 64-electrode cap with electrodes placed according to the international 10–10 system. Additionally, we placed two electrodes on the mastoids (roughly at TP9 and TP10). Data were pre-processed using EEGLAB^54^.

Signals were recorded at a 1024-Hz sampling rate, using the integrated CMS-DRL system as the online reference and later re-referenced offline to the average mastoid signal. Afterwards, the data were down-sampled to 256 Hz, and a high-pass filter of 0.1 Hz was applied. During manual inspection, we excluded bad channels (on average less than 2% of excluded channels) and blinks, eye movements and muscle activity were corrected by independent component analysis (ICA). In Experiment 1, triggers marking ping and search-array onset were shifted by 16 ms to correct for a systematic delay of one frame refresh between stimulus presentation on the screen and trigger transmission (measured by a photodiode), which was unnecessary for Experiment 2. Signals from excluded channels were then interpolated, and the continuous EEG data were segmented into epochs. For the analysis of anticipatory spatial biases, data were epoched from −200 to 600 ms relative to ping onset or, on no-ping trials, relative to the corresponding time point at which the ping would have appeared. To characterize neural activity during target selection and examine learning-related modulations of ERP components associated with selective and post-selective processing, search-locked epochs were extracted from −200 to 800 ms relative to search-array onset. Baseline correction was applied using the respective 200-ms pre-ping and pre-stimulus periods. Then, a second artefact rejection was performed to flag and remove epochs containing an absolute amplitude exceeding ±200 μV in the channels (on average excluding less than 5% of the trials).

### Pupillometry recording and pre-processing

Pupil data were recorded using an EyeLink 1000 eye tracker, sampling at 1000 Hz from either the left or right eye. A nine-point calibration was performed at the beginning of the experiment and repeated before each experimental block.

Data preprocessing was conducted with R (version 3.6.2)^55^. Missing pupil samples, as well as 80 ms before and after each gap, were interpolated using linear interpolation. When two such gaps occurred within 100 ms of each other, they were treated as a single gap and interpolated accordingly. Data were then epoched from −200 to 1900 ms relative to search array onset and baseline corrected using the -200 ms pre-stimulus period. Note that in the SL session of Experiment 2, in which no reward feedback was presented, the final 600 ms of the epoch overlapped with the initial portion of the subsequent trial (which represented no visible change, with just the fixation cross being visible). This was done to allow comparison with RL condition. Epochs with more than 50% of missing data were excluded (on average less than 5%). Finally, pupil data were downsampled to 250 Hz.

## Analysis

Statistical analyses were performed in R using the ez^56^ and effectsize^57^ packages. Behavioural analysis included the analysis of accuracy and RTs. For RTs, only correct responses were included. In all analyses, unless otherwise specified, the LFL and LRL conditions represent the average across the three low-frequency target/low-reward locations.

For five participants in Experiment 1 and for one participant in Experiment 2, eye-tracking data could not be recorded because the eye could not be reliably detected by the eyetracker. Furthermore, due to a technical issue, in the RL session, one participant had missing data from the last two blocks, and another participant had no EEG data for two blocks (but still have, and have used, behavioural and pupil data). Participants were therefore included in all analyses for which valid data were available, resulting in small differences in sample size across modalities and analyses.

### Decoding analysis

To determine whether RL spatial priorities could be decoded in a similar fashion to the one observed for SL before the search array onset (Experiment 1), and how they are compared to the one of SL and how they evolve over time after the search array (Experiment 2) we conducted MVPA using the Amsterdam Decoding and Modeling toolbox (ADAM)^58^.

The same general decoding procedure was applied across experiments, with analyses performed separately on ping-locked (Experiment 1 and 2) and search-array-locked EEG activity (Experiment 2). A Linear Discriminant Analysis (LDA) classifier was applied to activity recorded from all 64 scalp electrodes. To capture the representational dynamics at the moment of (possible) ping stimulation, decoding was performed from −200 to 600 ms relative to ping onset or, for no-ping trials in Experiment 1, relative to the corresponding time point at which the ping would otherwise have appeared. The end of the time window corresponded to the minimum time window available before presentation of the subsequent search display. For the search-array analysis, decoding was performed from −200 to 800 ms relative to search-array onset, encompassing the period of active target selection and response processing.

In Experiment 1, classifiers were trained to decode the learned HRL, with the four possible HRL positions (mid-left, mid-right, lower-left and lower-right) defining the four decoding classes. Decoding was performed separately for ping and no-ping trials, allowing us to assess whether reward-based spatial priority could be detected in anticipatory neural activity and whether its representation depended on perturbation by the ping stimulus.

A similar approach was used in Experiment 2 where decoding was performed separately for the SL and RL sessions. In the SL session, the four possible HFLs defined the decoding classes, whereas in the RL session the four possible HRLs served as class labels (as in Experiment 1). Here, the same four-class classification of the HFL and HRL were applied to decode activity elicited by the search-arrays. Note that, for search-array-locked decoding, the learned high-priority location defining a given class was independent of the physical location of the target on an individual trial. Thus, trials assigned to the same decoding class could contain targets at any of the four possible target locations. Above-chance classification therefore reflected neural information associated with the learned spatial priority rather than simply the target’s current physical position. However, to prevent decoding performance from being driven by imbalances in target location frequency or low-level target features (e.g., orientation or gap position), each decoding class was fully balanced to contain an equal number of trials for every actual target location and target feature combination. This balancing procedure was particularly critical in the SL session, in which targets appeared more frequently at one location relative to the other three locations.

To reduce the computational load, data were downsampled from 256 Hz to 128 Hz using ADAM’s resampling procedure, which applies an anti-aliasing low-pass filter before resampling. Classification performance was assessed using a 10-fold cross-validation procedure in which trials were randomly assigned to folds. At each time point, classifiers were trained on 90% of the data and evaluated on the remaining 10%, with this process being repeated until all folds had served as test data. Final decoding accuracy was computed as the average across folds. Classification performance was quantified using the area under the curve (AUC), with a value of 0.5 indicating chance-level decoding. At the group level, statistical significance of decoding performance for both SL and RL was assessed using cluster-based permutation tests (10,000 permutations) and paired sampled t-tests against chance. To compare decoding performance (Experiment 1: ping vs. no-ping trials; Experiment 2: SL vs RL), we used the same cluster-based permutation test approach (two-sided, 10,000 permutations).

### Pupillometry analysis

To examine whether task-evoked pupil responses reflected attentional selection driven by RL, and subsequently to compare these effects with SL, we quantified mean baseline-corrected pupil diameter between 800 and 1900 ms following search-array onset. The start of this window was chosen in line with the known latency of cognitive effects on pupil size that peaks around 1 s following stimulus onset^59^. The end of the window was chosen to provide a common analysis interval across learning conditions while accounting for differences in trial duration. Specifically, because RL trials included an additional reward-feedback period, they were longer than SL trials. Importantly, although the analysis window extended into the reward- feedback period in RL trials, feedback-related effects are unlikely to substantially contribute because of the slow dynamics of pupil responses and the fact that visual feedback was highly visually similar for high- and low-reward conditions. The resulting values were analysed using a rm-ANOVA, with experimental condition as a within-subject factor. Only correct trials were included in these analyses.

In addition, in Experiment 2, absolute pupil size was analysed to assess whether the RL and SL sessions were generally approached with a similar level of arousal/effort by the participants on a block-level. To this end, we computed, for each subject, the mean pupil diameter (without baseline correction) within a neutral time window preceding the onset of the ping display and compared these values across experimental sessions. Specifically, the time window selected was from -200 ms to the ping display onset.

### Computational modelling

To link learning dynamics with decision-making, behavioural data collected in Experiment 2 were modelled using a combination of the RW^35^ learning rule and the hierarchical Bayesian methods to estimate parameters in the DDM^36^. Trials with RTs faster than 200 ms were excluded and analysis was implemented in Python using the HDDM (v 0.6) package^60^. The RW model was first used to estimate trial-by-trial Q-values separately for the SL and RL sessions, and these estimates were subsequently incorporated into the HDDM as trial-wise regressors of decision parameters. For both SL and RL session, the expected value (Q-value) associated with each of the four possible target locations was initialized uniformly (Q = 0.25), reflecting the absence of prior knowledge about the spatial regularities. Q-values were updated on each trial according to the Rescorla–Wagner learning rule:

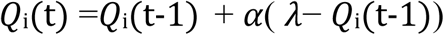

where *Q*i(t) represents the expected value associated with location *i* at trial *t*, *Q*i(t-1) is the value from the previous trial, *λ* denotes the outcome, and *α* is the learning rate. In the SL session, *λ* was coded as 1 for the observed target location (i.e., where the target appeared) and 0 for all non-observed locations. In the RL session, when response was correct, *λ* was coded as 1 if target appeared at HRL or 0.001 if target appeared at LRL. For the remaining three locations *λ* was 0. When a response was incorrect *λ* was also coded as 0 for all the four locations. We estimated learning rate via a grid search over 13 equally spaced learning-rate values in the range 0.001-0.8. To provide a baseline model without learning, we instead estimated HDDMs that did not include the RW-derived learning regressor (or equivalently, with *α* = 0).

For each trial, a single Q-difference was derived by computing the difference between the Q-value of the observed target location and the mean Q-value of the remaining locations:

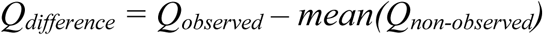

This trial-wise Q-difference signal quantified the relative advantage of the current target location over the alternative locations. Consequently, higher values indicated a stronger learned preference for the HFL in the SL session or the HRL in the RL session, whereas lower values reflected weaker learned preferences for LFL and LRL.

The trial-wise Q-difference signal was then entered as a parametric regressor in a HDDM. Separate regression models were estimated in which the Q-difference signal modulated drift rate (v), boundary separation (a), non-decision time (t), or starting point (z) individually, together with a full model in which the Q-difference signal simultaneously modulated all four parameters. Models were estimated using Markov chain Monte Carlo (MCMC) sampling with four chains of 2,000 samples each, with the first 500 samples discarded as burn-in to ensure convergence and accurate posterior probability estimation. Model comparison was performed using the DIC, with lower values indicating a better model fit^61^. To assess model convergence, the Gelman-Rubin R-hat statistic was computed^62^. Results indicated that all parameters yielding R-hat values below 1.1^63^.

### Event-related potentials (ERPs) analysis

ERP analyses were performed using ERPLAB^64^. The ERP markers of interest were the N2pc and the SPCN. These components were quantified at posterior–occipital electrodes PO7 and PO8, where their amplitudes are known to be maximal^26,65^. Based on established conventions in the literature, the N2pc was measured within a 200–300 ms interval following search array onset^25,66^, whereas the SPCN was examined in a later window spanning 350–650 ms post-onset^67,68^.

For each participant, mean amplitudes were extracted separately for electrodes contralateral and ipsilateral to the target position and then averaged across the experimental blocks. Lateralized activity was computed by subtracting ipsilateral from contralateral waveforms, and these difference scores served as dependent measures in rm-ANOVA assessing effects across experimental conditions. Only correct trials were included in these analyses.

## Data availability

The anonymized behavioural, EEG and eye-tracking data generated in this study will be available on EBRAINS repository after publication.

## Code availability

Scripts related to behavioural, EEG and eye-tracking preprocessing and main analysis, will be available on EBRAINS repository after publication.

## Supporting information

Supplementary Analyses of the main manuscript

