## Supplementary Analyses of the main manuscript for "Common attentional mechanisms, but distinct computational dynamics, across Statistical and Reward Learning"

**ERP results of Experiment 1**

In addition to the latent neural activity, EEG data elicited by the onset of the search array were used to examine two well-established ERP components indexing attentional selection and post-selection processing: the N2pc and the SPCN, respectively. This analysis aimed to further characterize the involvement of RL during spatial attention.

In terms of the N2pc, we conducted a one-way rm-ANOVA with Reward Location (high, low) on the time-window of 200-300 ms after stimulus onset. However, this analysis revealed no significant main effects of Reward Location [$F(1,44) = 0.93$, $p= 0.33$].

To investigate later stages of target processing, we analysed the SPCN in the 350–650 ms time window using the same one-way rm-ANOVA with Reward Location (high, low). Also in this case, no main effect was observed for Reward Location [$F(1,44) = 0.14$, $p = 0.70$]. Importantly, the absence of significant ERP effects does not necessarily indicate that RL failed to influence spatial attention, but rather that ERPs capture only specific stimulus-locked components of neural processing and may therefore be insensitive to distributed latent changes in spatial priority detectable with multivariate approaches (for this, see main text).

**ERP results of Experiment 2**

In Experiment 2 we wanted to compare RL and SL in terms of N2pc and SPCN. As in Experiment 1, to analyse the N2pc, we conducted a 2×2 rm-ANOVA with Learning Mechanism (SL, RL) and Location (high, low) on the time-window of 200-300 ms after stimulus onset. This analysis revealed no significant main effects of Learning Mechanism [$F(1,29) < 0.01$, $p= 0.98$] or Location [$F(1,29) = 0.06$, $p= 0.79$]. The interaction between the two factor was also not significant [$F(1,29) = 1.83$, $p= 0.18$] (Fig. 5a).

To investigate further target processing, we analysed the SPCN in the 350–650 ms time window using the same 2×2 rm-ANOVA with Learning Mechanism (SL, RL) and Location (high, low). Also in this case, no main effect was observed for Learning Mechanism [$F(1,29) = 2.49$, $p = 0.12$] or Location [$F(1,29) = 1.69$, $p = 0.20$]. However, a trend to significant interaction emerged [$F(1,29) = 2.98$, $p= 0.09$, $\eta^2_p = 0.09$]. Interestingly, planned post-hoc t-test (two-tailed) showed a significant difference between the high (vs. low) Location, during the RL-session [$t(29) = 2.66$; $p = 0.01$, Cohen's $d = 0.35$] but not during SL-session [$t(29) = 0.15$; $p = 0.87$] (Fig. 5b).

Although the N2pc did not reveal attentional modulation by either learning mechanism, the (post-hoc) SPCN results suggest that RL may selectively modulate later stages of target selection. Specifically, RL may reflect increased post-selection processing demands for locations with lower attentional priority. Yet, it is important to note that Experiment 1 did not show such an effect.

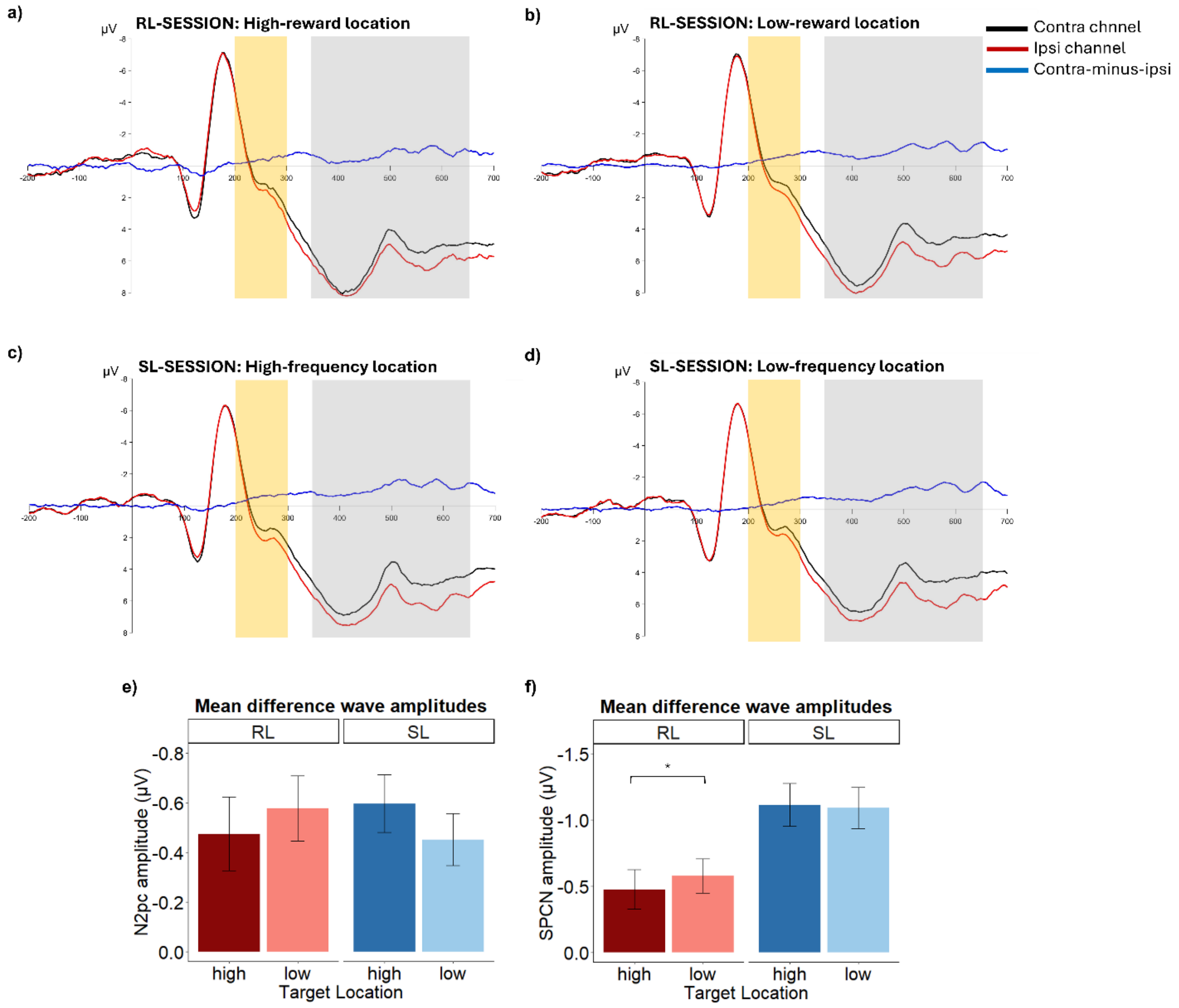

**Figure 5 | N2pc and SPCN as a function of learning mechanism and target location**. Sensor waveforms showing contralateral (black), ipsilateral (red), and difference (contra − ipsi; blue) activity relative to the target during the RL (**a, b**) and SL (**c, d**) sessions. Panels **a** and **c** show waveforms for high-reward and high-frequency target locations, respectively, whereas panels **b** and **d** show waveforms for low-reward and low-frequency target locations. Time 0 ms corresponds to search-display onset. The yellow area indicates the N2pc measurement window (200–300 ms), and the gray area indicates the SPCN measurement window (350–650 ms). **e**, Mean N2pc amplitude (contralateral − ipsilateral) for high and low locations as a function of learning mechanism (RL vs. SL). **f**, Mean SPCN amplitude (350–650 ms; contralateral − ipsilateral) for high and low locations as a function of learning mechanism (RL vs. SL). Error bars represent s.e.m. Asterisks indicate significant pairwise comparisons ($p < 0.05$).

**Inter-trial priming effects do not compromise decoding results of Experiment 2**

To assess whether the ping-evoked decoding observed following SL could be explained by inter-trial priming we performed a control analysis in which the frequency of target locations on the preceding trial was equated across conditions. Specifically, we matched the number of trials in which the target had previously appeared at the HFL/HRL and at each of the three remaining LFL/LRL. Note that, unlike in the control behavioural analysis, here we did not exclude immediate target location repetition, because the target had not yet appeared at the time of ping onset. As shown in Figure 6, following this exclusion, decoding in the SL session remained significantly above chance, even if less sustained (119-307 ms). This demonstrated that anticipatory neural representations associated with SL cannot be explained solely by short-term effects of HFL, but rather, that it reflects a long-lasting and stable attentional bias. Likewise, decoding in the RL session also remained above chance (135-236 ms and 268-322 ms). Importantly, the direct comparison between learning conditions remained non-significant, confirming a similarity between the two sources of learning.

Also for the EEG activity evoked by search array, as a control analysis we checked whether the decoding observed especially in the SL session could be explained by short-term inter-trial priming. We thus repeated the analysis after excluding trials in which the target appeared at the same location as on the preceding trial. Unlike the ping-decoding analysis, the identity of the current target location was known during the search display, allowing consecutive target repetitions to be removed directly, consistent with the behavioural control analyses. Despite this control, classifier performance during SL session remained reliably above chance (139-732 ms). RL also confirmed sustained above chance decoding performance (170-537 ms and 553-709 ms). Notably, the similarity between the primary (with no exclusion of consecutive target-location repetition) and this control analyses likely reflects the balancing procedure implemented in the main decoding analysis, which already removed a substantial proportion of consecutive repetitions while matching target locations within each classification labels. Moreover, a two-tailed cluster-based permutation t-test again revealed no significant differences between the SL and RL sessions, further supporting convergence during active visual search.

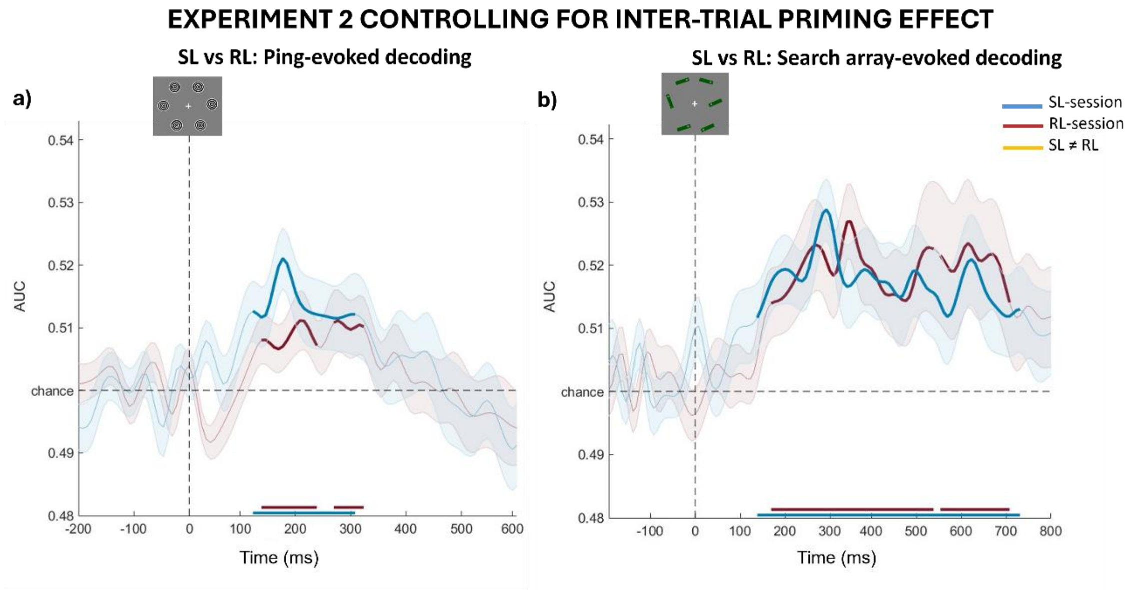


**Figure 6 | Decoding of SL and RL after controlling for inter-trial priming a,** Decoding accuracy time-locked to the onset of the task-irrelevant ping display and **b,** search array, shown separately for SL (blue) and RL (red). In all panels, chance-level decoding corresponds to 0.5. Horizontal bars above the x-axis indicate significant temporal clusters identified using cluster-based permutation tests ($p < 0.05$) against chance. In neither case did we observe any differences between SL and RL (as indicated by the absence of yellow lines marking significant temporal clusters). Shaded areas represent the s.e.m. across participants. Curves were smoothed for visualization using a 37-Hz low-pass filter.

Furthermore, we also controlled for inter-trial priming effect on pupil size. Specifically, we replicated the 2×2 rm-ANOVA with Learning Paradigm (SL, RL) and Location (high, low), but excluding trials in which the target appeared in the same spatial location as in the previous trial. Results confirmed a significant main effect of Learning Paradigm [$F(1,28) = 29.06$, $p < 0.01$, $\eta^2_p = 0.50$], and a significant interaction between the two factors [$F(1,28) = 4.42$, $p = 0.04$, $\eta^2_p = 0.13$], while again not finding significant main effect of Location [$F(1,28) = 0.09$, $p = 0.76$]. Post-hoc t-test (two-tailed) did not show any significant differences between the high (vs. low) Location, neither during the SL session [$t(28) = 1.54$; $p = 0.13$] nor during RL session [$t(28) = 1.57$; $p = 0.12$], suggesting that the interaction arose from a mixture of these two effects (numerically larger for LFL than HFL and for HRL than LRL).

Finally, turning to computational modelling, in the SL session we controlled for inter-trial priming effect by entering the Q-difference into HDDM after removing trials where target appeared in the same location as in the previous trial. Even after this exclusion of trials, results revealed higher Q-difference values were still associated with a significant increase in drift rate (v; posterior mean = 0.41, 95% credible interval (CrI) = [0.20, 0.58]), confirming faster accumulation of sensory evidence for targets appearing at the learned HFL. However, we did not replicate evidence in terms of non-decision time (t; posterior mean = −0.004, 95% CrI = [−0.04, 0.03]). Starting point (z; posterior mean = 0.003, 95% CrI = [−0.02, 0.02]) and boundary separation (a; posterior mean = −0.05, 95% CrI = [−0.15, 0.04]) remained non-significant.

**Alternative model for RL tested in Experiment 2**

The learning algorithm described in the main text assumes that value estimates are updated for all locations on every trial. This assumption is intuitive for SL, where the occurrence of a target at one location simultaneously provides information about the remaining locations (where hence the target did not appear on this trial). In RL, however, such an assumption is less straightforward because reward outcomes are contingent on the selected target location and do not necessarily imply a reduction in the value of unselected locations.

We therefore tested an alternative, conditional RL model in which, for each trial, only the value associated with the target location was updated. Specifically, when a target appeared at a given location, its expected value (Q-value) was updated according to the same Rescorla-Wagner learning rule described in the main text, whereas the values associated with all other locations remained unchanged. Model comparison based on the DIC indicated that this alternative model provided a poorer fit to the data than the model reported in the main text.
